# AVE0991, a Mas receptor agonist, increases influenza and COVID-19 severity *in vivo*

**DOI:** 10.64898/2026.02.23.707600

**Authors:** Melanie Wu, Julian D.J. Sng, Ellesandra C. Noye, Georgina McCallum, Helle Bielefeldt-Ohmann, Cheng Xiang Foo, Katharina Ronacher, Keng Yih Chew, Kirsty R. Short

**Author notes:** **Corresponding author:** Kirsty Short, School of Chemistry and Molecular Biosciences, The University of Queensland, Brisbane, QLD 4072, Australia.

## Abstract

Dysregulation of the renin-angiotensin system (RAS) contributes to severe influenza and COVID-19, potentially via impaired ACE2/Ang-(1-7)/Mas receptor (MasR) signalling. AVE0991, an orally bioavailable MasR agonist, protects against non-infectious lung inflammation, but its effects in viral respiratory disease are unknown. We evaluated AVE0991 in human lung epithelial cells and murine models of influenza A virus (IAV) and SARS-CoV-2 infection. *In vitro*, AVE0991 suppressed cytokines IL-6 and TNF-α to both IAV and SARS-CoV-2 and reduced IAV titres. Unexpectedly, *in vivo* treatment worsened disease. These findings highlight discordant *in vitro* and *in vivo* effects and underscore the need for careful evaluation of RAS-targeted therapies in acute viral infection.

## Introduction

Severe influenza and COVID-19 are driven by dysregulated immune responses that contribute to tissue damage and acute respiratory distress syndrome (ARDS) [1]. The renin-angiotensin system (RAS) has gained attention as a therapeutic target, as SARS-CoV-2 enters cells via angiotensin-converting enzyme 2 (ACE2), which degrades the pro-inflammatory mediator angiotensin II (Ang II) [2]. ACE2 internalization during infection is thought to disrupt RAS balance, as reflected by altered RAS metabolites in severe COVID-19 [2].

Although influenza does not engage ACE2 for entry, severe infections have similarly been associated with RAS imbalance. Protective RAS signalling, mediated ACE2/Angiotensin-(1-7) [Ang-(1-7)]/Mas receptor (MasR) signalling, has been linked to reduced inflammation and improved pulmonary outcomes in severe influenza [3]. Recombinant Ang-(1-7) protects in preclinical models of ARDS but shows limited clinical translation [4, 5]. Ang-(1-7) also has pharmacokinetic limitations, including a short half-life and the need for intravenous administration. In contrast, AVE0991, a non-peptide analogue of Ang-(1-7), is orally bioavailable, resistant to enzymatic degradation, has a longer half-life, and selectively activates MasR to mimic Ang-(1-7)’s protective effects [6].

Although AVE0991 shows promise in non-infectious models of chronic inflammation and injury [7], and modulators of RAS have been widely stipulated to reduce viral disease [3, 5]. AVE0991 has yet to be tested in acute models of respiratory viral infection. In this study, we assessed its therapeutic potential using both *in vitro* and *in vivo* models of SARS-CoV-2 and influenza virus infection.

## Materials & Methods

### Ethics

All experiments were approved by the University of Queensland Animal Ethics Committee (2024/AE000314, 2021/AE000674, and 2021/AE000089).

### Viruses

IAV (A/H1N1/Auckland/1/2009) was propagated in embryonated eggs as previously described [8]. Delta SARS-CoV-2 strain (EPI_ISL_2433928) was used *in vitro* and was provided by Queensland Department of Health. A mouse-adapted SARS-CoV-2 strain (EPI_ISL_968081) generated in Foo et. al [9], was used for *in vivo* studies. Viral titres were determined by plaque assays as detailed in the Supplementary Material.

### Cell infection models

Calu-3 cells (ATCC; HTB-55) were maintained as described in Supplementary Material. Cells were infected with IAV (MOI 0.1) or Delta SARS-CoV-2 (MOI 0.01) for 1 h, washed with PBS, and cultured for 24 h. AVE0991 or vehicle (0.125% DMSO) was added at 24 h post-infection. At 48 h, supernatants were collected for viral titres, and cell lysates harvested for RNA extraction and qPCR.

### Mouse models

Male C57BL/6J mice (13 weeks for IAV; 20 weeks for SARS-CoV-2) were intranasally infected with IAV (2,500 PFU) or mouse-adapted SARS-CoV-2 (10^4^ PFU) under anaesthesia. AVE0991 (10 mg·kg^−1^) or vehicle was administered twice daily by oral gavage starting at 2 dpi (IAV) or 1.5 dpi (SARS-CoV-2) until endpoint. Blood oxygen saturation was measured in IAV-infected mice at endpoint as described in Supplementary Material. Animals were euthanized and tissues collected for downstream analyses detailed in Supplementary Material.

### RNA extraction and qPCR

Total RNA was extracted, reverse transcribed, and analysed by SYBR-based qPCR using the ΔΔCt method normalized to GAPDH. Detailed information is in the Supplemental Material. Primer sequences are provided in Supplemental Table 1.

### Flow cytometry

Flow cytometry was performed on single-cell suspensions from enzymatically digested lungs, as detailed in the supplementary material, and analysed using the gating strategy shown in Supplemental Figure 1.

### Histological Analysis

The left lung lobe was fixed in 10% neutral-buffered formalin for ≥24 hours, then transferred to 70% ethanol for processing and haematoxylin and eosin (H&E) staining at the Core Histology Facility, Translational Research Institute. Sections were scored by a blinded veterinary pathologist (see Supplementary Material).

### Multiplex cytokine assay

Serum cytokines were quantified using a multiplex bead-based assay (Supplementary Material).

### Statistical analyses

Data were analysed using GraphPad Prism v10.6. Outliers were identified using ROUT (Q=1%). Normality was assessed by Shapiro–Wilk testing. Two-group comparisons used unpaired t-tests or Mann–Whitney tests as appropriate. Weight loss was analysed by two-way ANOVA with Geisser–Greenhouse correction and Tukey post-hoc testing.

## Results

### AVE0991 reduces cytokine gene expression during IAV and SARS-CoV-2 infection and reduces IAV titres *in vitro*

To investigate whether AVE0991 modulates cytokine production, a hallmark feature of severe viral infection, we used Calu-3 lung epithelial cells to model treatment *in vitro*. Toxicity assays showed AVE0991 is cytotoxic in Calu-3 cells at concentrations ≥25 μM (Supplemental Figure 2); therefore, a concentration of 12.5 μM was used for all subsequent experiments. Calu-3 cells were infected with either IAV or Delta variant SARS-CoV-2 for 24 h, followed by treatment with AVE0991 (12.5 μM) for an additional 24 h. In both infection models, AVE0991 significantly reduced expression of the pro-inflammatory cytokines IL-6 and TNF-α (Figures 1A, 1C). Interestingly, while AVE0991 reduced IAV titres (Figure 1B), it had no impact on SARS-CoV-2 titres (Figure 1D). These findings suggest that AVE0991 exerts anti-inflammatory effects in response to both viruses and may have antiviral activity specifically against IAV.

**Figure 1.**
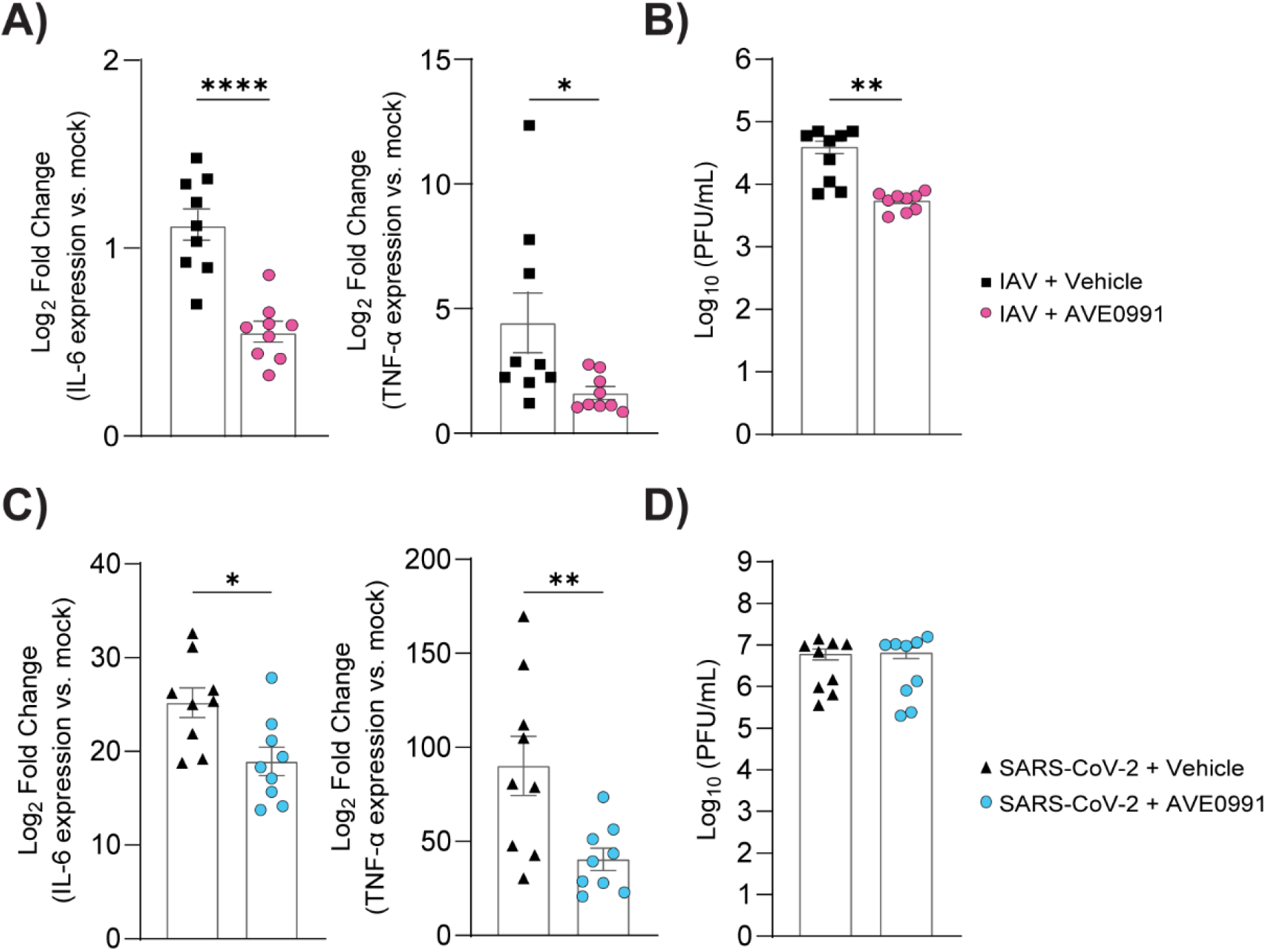
AVE0991 reduces cytokine gene expression during IAV and SARS-CoV-2 infection. Calu-3 cells were infected with MOI 0.1 IAV (A, B) or MOI 0.01 SARS-CoV-2 Delta (C, D) for 24 hours, then treated with vehicle (0.125% DMSO) or AVE0991 (12.5 μM) for 24 h. (A, C) Log_2_ fold change in IL-6 and TNF-α gene expression. (B, D) Viral titres in supernatants (PFU/mL) measured by plaque assay. Normality was assessed by Shapiro-Wilk test. Statistical significance was determined using unpaired t-test or Mann-Whitney test, as appropriate. Data represents three independent replicates and are shown as mean ± SEM. *: p<0.05; **: p<0.01; ****: p<0.0001.

### AVE0991 increased IAV and SARS-CoV-2 disease severity *in vivo*

Building on promising *in vitro* results, AVE0991 was next evaluated *in vivo* using established mouse models of IAV and SARS-CoV-2 infection [9]. A dose of 10 mg/kg, administered twice daily via oral gavage, was confirmed non-toxic based on body weight, pulse oximetry, and serum biomarkers (Supplemental Figure 3). Disease severity was assessed by monitoring weight loss, the standard endpoint in influenza models [9], and blood oxygen saturation for IAV, a direct indicator of lung function [10]. Compared to vehicle controls, AVE0991-treated mice experienced significantly greater weight loss in both infection models (Figure 2A). Blood oxygen saturation remained unchanged in IAV-infected mice (Supplemental Figure 4). Lung histology revealed a trend toward increased tissue damage in IAV + AVE0991 mice (p = 0.0558), while no histological differences were observed in SARS-CoV-2-infected mice (Supplemental Figure 4). Viral titres were elevated in AVE0991-treated IAV mice but remained unchanged in SARS-CoV-2 groups (Figure 2B). Immune profiling showed a significant reduction in alveolar macrophages in IAV + AVE0991 mice, suggesting compromised protection in the treated group [9] (Figure 2C). In contrast, T cell frequencies were increased, though the functional implications remain unclear. No significant changes in immune cell populations were detected in SARS-CoV-2-infected mice (Figure 2C). Serum cytokine analysis showed reduced IFNγ in the IAV + AVE0991 group (Figure 2D), while CXCL1 and CXCL10 levels were elevated in SARS-CoV-2 + AVE0991 mice. No other cytokines were significantly altered (Supplemental Figure 5). Overall, these findings suggest that although AVE0991 modulates immune responses during infection, this may exacerbate disease severity in both IAV and SARS-CoV-2 mouse models.

**Figure 2.**
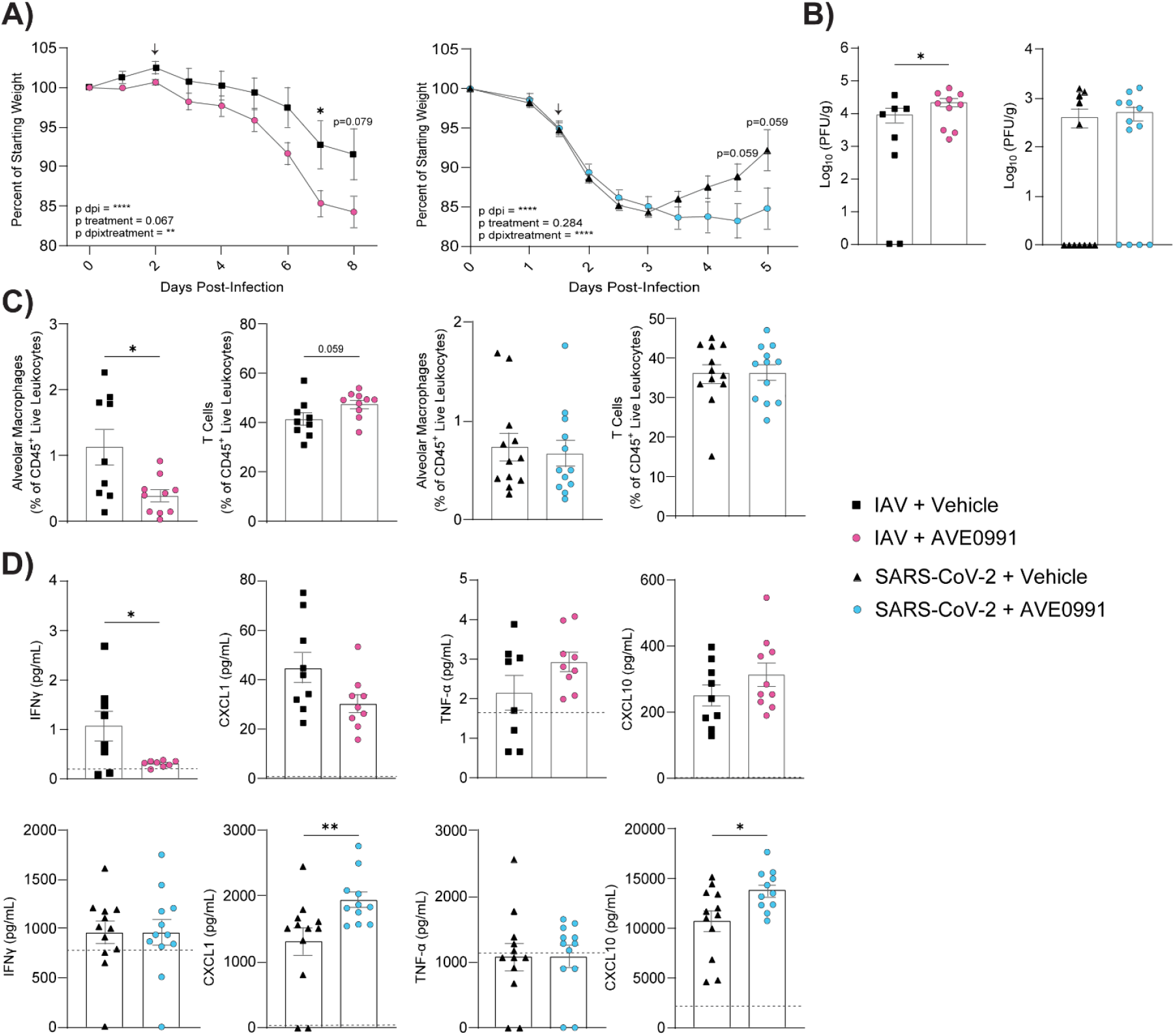
AVE0991 increases IAV and SARS-CoV-2 disease severity in vivo. C57BL/6J mice (13 or 20 weeks old) were infected intranasally with 2,500 PFU of IAV or 10^4^ PFU of SARS-CoV-2, respectively. Mice received AVE0991 (10 mg/kg) or vehicle via oral gavage twice daily (12-hour intervals), starting at 2 dpi (IAV) or 1.5 dpi (SARS-CoV-2) until the endpoint. (A) Body weight (expressed as % of starting weight). Arrow indicates treatment initiation. (B) Viral titres (PFU/g) in lung homogenates. (C) Frequencies of immune cell subsets (% of CD45^+^ live leukocytes) from lungs. (D) Serum cytokine levels (pg/mL); dotted line indicates the limit of detection of the assay. Statistical analysis: (A) two-way ANOVA with Geisser-Greenhouse correction and Tukey’s multiple comparisons test; (B–D) each point represents an individual mouse, outliers were removed using ROUT (Q = 1%), and unpaired t-test or Mann–Whitney test was applied as appropriate. Data are pooled from two independent experiments and presented as mean ± SEM. *: p<0.05; **: p<0.01; ****: p<0.0001.

## Discussion

Dysregulation of the RAS is a hallmark of severe disease in IAV and SARS-CoV-2 infections and enhancing the ACE2/Ang-(1-7)/MasR axis has been proposed as a therapeutic strategy [2, 3]. Recombinant Ang-(1-7) improves outcomes in experimental models of viral respiratory disease [11], and the synthetic MasR agonist AVE0991, offering improved stability and oral bioavailability, has demonstrated protective effects in models of non-infectious lung injury [6, 7]. However, its efficacy during viral infection remains unknown. Here, we show that although AVE0991 reduced pro-inflammatory cytokine production in infected epithelial cells *in vitro*, it unexpectedly worsened disease *in vivo*, highlighting a context-dependent role for MasR activation in viral pathogenesis.

AVE0991 suppressed IL-6 and TNF-α expression in IAV- and SARS-CoV-2-infected Calu-3 cells, extending its known anti-inflammatory effects to virally infected lung epithelium [12]. It also reduced IAV titres *in vitro*, suggesting a potential influenza-specific antiviral effect, although no antiviral activity was observed for SARS-CoV-2. In contrast, in our *in vivo* IAV experiments, AVE0991 treatment did not alter viral load. A prior study reported that Ang-(1– 7) reduced viral titres in IAV-infected mice [11], but differences in compound, dosing, and timing likely contribute to these divergent outcomes. Additional distinctions may also reflect the broader receptor activity of Ang-(1–7), including signalling via MrgD and AT_2_R [13].

Despite these *in vitro* effects, AVE0991 worsened disease in both infection models, with increased weight loss and virus-specific cytokine alterations, including reduced IFN-γ in IAV and elevated CXCL1/CXCL10 in SARS-CoV-2. During acute viral infection, early pro-inflammatory and interferon responses are critical for viral clearance [14]. Premature suppression of these pathways may impair host defence. In contrast, AVE0991 is beneficial in chronic inflammatory models [7], suggesting that disease context and timing critically determine therapeutic outcomes. Parallels can also be drawn with corticosteroid treatment in severe IAV and COVID-19, where early administration impaired host defence and worsened outcomes, while delayed treatment in severe COVID-19 improved survival [15]. As treatment in this study commenced at 1.5–2 dpi, future work should assess whether delayed administration alters disease severity and define the mechanisms underlying MasR signalling *in vivo*.

In summary, these *in vivo* findings demonstrate that MasR activation can have deleterious effects during acute viral infection, despite its anti-inflammatory activity *in vitro*. Our results suggest that contrary to the conventional view [5], MasR activation may have context-dependent effects during respiratory viral infections. Although this pathway is generally considered anti-inflammatory, elevated levels of Ang-(1-7) and soluble ACE2 have been reported in patients with severe COVID-19 [2], highlighting the need for a better understanding of protective RAS signalling. Untimely MasR activation may disrupt immune homeostasis required for viral clearance. Further research is needed to clarify kinetics of MasR activation on disease severity, including studies on timing, dosing, and the potential use of MasR antagonists. Overall, the worsening of disease with AVE0991 in murine viral respiratory infection models highlights the need for context-specific evaluation when repurposing RAS-targeting therapies.

