## Supplemental_Material for "AVE0991, a Mas receptor agonist, increases influenza and COVID-19 severity *in vivo*"

### Supplemental Methods

*Cell Culture Maintenance*

Calu-3 cells (ATCC; HTB-55) were maintained in Minimum Essential Medium (MEM; Gibco) supplemented with 10% fetal calf serum (FCS; Sigma-Aldrich), 1% penicillin-streptomycin (Gibco), and 1% GlutaMAX (Gibco).

*Mouse Model*

C57BL/6J mice were obtained from Ozgene ARC and housed in individually ventilated cages under 12-hour light/dark cycles, with *ad libitum* access to food and water.

*Quantification of IAV Titres*

IAV plaque assays were performed on confluent monolayers of MDCK cells (ATCC NBL-2) using a semi-solid overlay. MDCK cells were maintained in Dulbecco's Modified Eagle Medium (DMEM; Gibco) supplemented with 10% fetal bovine serum (FBS; Gibco) and 1% penicillin-streptomycin (Gibco). Cells were seeded into 12-well plates at a density of 700,000 cells per well. The following day, cells were washed once with serum-free DMEM and inoculated with 10-fold serial dilutions of virus stocks or experimental samples. Plates were incubated at 37 °C with 5% $CO_2$ for 1 hour, with rocking every 15 minutes. After incubation, wells were overlaid with a semi-solid medium consisting of serum-free MEM (Gibco) containing 0.1% low EEO agarose (Sigma-Aldrich) and 1 μg/mL TPCK-treated trypsin (Worthington Biochemical). Plates were incubated for 3 days, after which the overlay was removed, and cells were stained with crystal violet solution (0.1% crystal violet in 10% neutral buffered formalin) for 1 hour to visualize plaques.

*Quantification of SARS-CoV-2 Titres*

SARS-CoV-2 plaque assays were performed on confluent monolayers of Vero E6 (ATCC CRL-1586) cells using a semi-solid overlay. Vero E6 cells were maintained in DMEM (Gibco) supplemented with 10% FBS and 1% penicillin-streptomycin (Gibco). Cells were seeded into 12-well plates at a density of 200,000 cells per well. After two days, cells were inoculated with 10-fold serial dilutions of relevant virus stocks or experimental samples. Plates were incubated at 37 °C with 5% $CO_2$ for 1 hour, with rocking every 15 minutes. After incubation, wells were overlaid with semi-solid medium consisting of MEM (Gibco) containing 2% FBS and 0.1% low EEO agarose (Sigma-Aldrich). Plates were incubated for 2 days (for mouse-adapted SARS-CoV-2) or 3 days (for Delta SARS-CoV-2), after which the overlay was removed, and cells were stained with crystal violet solution (0.1% crystal violet in 10% neutral buffered formalin) for 1 hour to visualize plaques.

*RNA extraction and reverse-transcriptase quantitative PCR (qPCR)*

Total RNA was extracted using the NucleoZOL kit (Takara Bioscience) and reverse transcribed with the High-Capacity cDNA Reverse Transcription Kit (Applied Biosystems). qPCR was performed on cDNA using SYBR Green (Invitrogen) on a QuantStudio 6 Flex system (Applied Biosystems), following manufacturer instructions.

*Cell Viability Assay*

Calu-3 cells were seeded at a density of 250,000 cells per well in black-walled 96-well plates. The following day, cells were washed once with PBS and treated with various concentrations of AVE0991 (6.25 μM, 12.5 μM, or 25 μM), vehicle control (DMSO in serum-free media), or media-only controls. Cell viability was assessed 24 hours post-treatment using the ATPlite 1-Step Luminescence Assay Kit (PerkinElmer) according to the manufacturer's instructions. Luminescence was measured using a CLARIOstar Plus plate reader.

*Mouse Drug Dosing*

13-week-old mice were treated with 10 mg/kg AVE0991 twice daily at 12-hour intervals for four consecutive days. At four days post-treatment, blood oxygen saturation was measured using the MouseOx pulse oximeter with a sensor collar. Mice were then euthanised, and blood was collected via cardiac puncture for downstream analysis of toxicity biomarkers.

*Mouse Blood Processing*

After collection, blood samples were stored overnight at 4°C and serum was procured by centrifugation of whole-blood samples at 10,000g for 10 minutes.

*Mouse Blood Toxicity Detection*

Serum biochemical profiling for liver toxicity (ALT and AST) was performed at the Veterinary Laboratory Services, The University of Queensland, using OSR6109 (AST detection, Beckman

Coulter) and OSR6107 (ALT detection, Beckman Coulter) reagents on a Beckman Coulter AU480 analyser.

*Mouse Lung Homogenization*

The right superior, middle, inferior, and post-caval lung lobes were harvested into DMEM (Gibco) and mechanically disrupted using a Qiagen Tissuelyser II (Qiagen).

*Mouse Lung Digestion*

Lungs were harvested from C57BL/6 mice into serum-free DMEM and transferred to 1 mL of digestion medium containing 0.25 mg/mL Liberase™ (Roche) and 0.15 mg/mL DNase I (Thermo Fisher Scientific) in DMEM (Gibco). The tissue was minced with scissors and incubated at 37 °C for 20 minutes. The digested sample was then filtered through a 70 μm cell strainer into a fresh centrifuge tube. Cells were pelleted by centrifugation at 400 x g for 5 minutes and resuspended in 1 mL of RBC lysis buffer (Invitrogen) for 1 minute at 25 °C. The suspension was then diluted with 25 mL PBS, centrifuged again, and finally resuspended in 100 μL PBS containing EDTA (Sigma-Aldrich) for flow cytometry staining.

*Flow Cytometry Staining*

Single-cell suspensions were centrifuged at 400 x g for 5 minutes in 96-well round-bottom plates (Corning) and incubated for 20 minutes with Fixable Viability Stain 700 (1:15,000; 564997, BD Biosciences). Lung samples were then incubated with 50 μL of Fc Block Rat Anti-mouse (CD16/32, BD Biosciences), followed by staining with the antibody panel listed in Supplemental Table 2. Single-color compensation controls were prepared using either remaining mouse lung cells or VersaComp Antibody Capture Kit (Beckman Coulter). Both compensation controls and samples were washed twice with 200 μL PBS, fixed in 100 μL BD Cytofix Fixation Buffer (BD Biosciences), washed again, and transferred to 5 mL FACS tubes (Corning) for acquisition on a BD LSRFortessa X20 at the Translational Research Institute. Flow cytometry data were analysed using FlowJo v10.8 (Windows), with the gating strategy shown in Supplemental Figure 1.

*Histology*

After euthanasia, murine lungs were inflated via intratracheal administration of PBS. The left lung lobe was fixed in 10% neutral-buffered formalin for at least 24 hours before being transferred to 70% ethanol for routine processing at the Core Histology Facility, Translational Research Institute. A veterinary pathologist, blinded to the study design, scored the sections. The assessment included vascular changes, bronchitis, interstitial inflammation, alveolar inflammation, pneumocyte hypertrophy, and pleuritis. The term "lung parenchyma changes" denotes the combined scores for bronchitis, interstitial inflammation, alveolar inflammation, pneumocyte hypertrophy, and pleuritis.

*Multiplex cytokine assay*

A panel of 13 cytokines, including IFN-γ, CXCL1, TNF-α, CCL2, IL-12p70, CCL5, IL-1β, CXCL10, GM-CSF, IL-10, IFN-β, IFN-α, and IL-6, was measured in mouse serum using the LEGENDplex Mouse Anti-Virus Response Panel (BioLegend) following manufacturer's instructions. Samples were run on a BD Accuri C6 Plus flow cytometer (BD Biosciences).

### Supplemental Figures

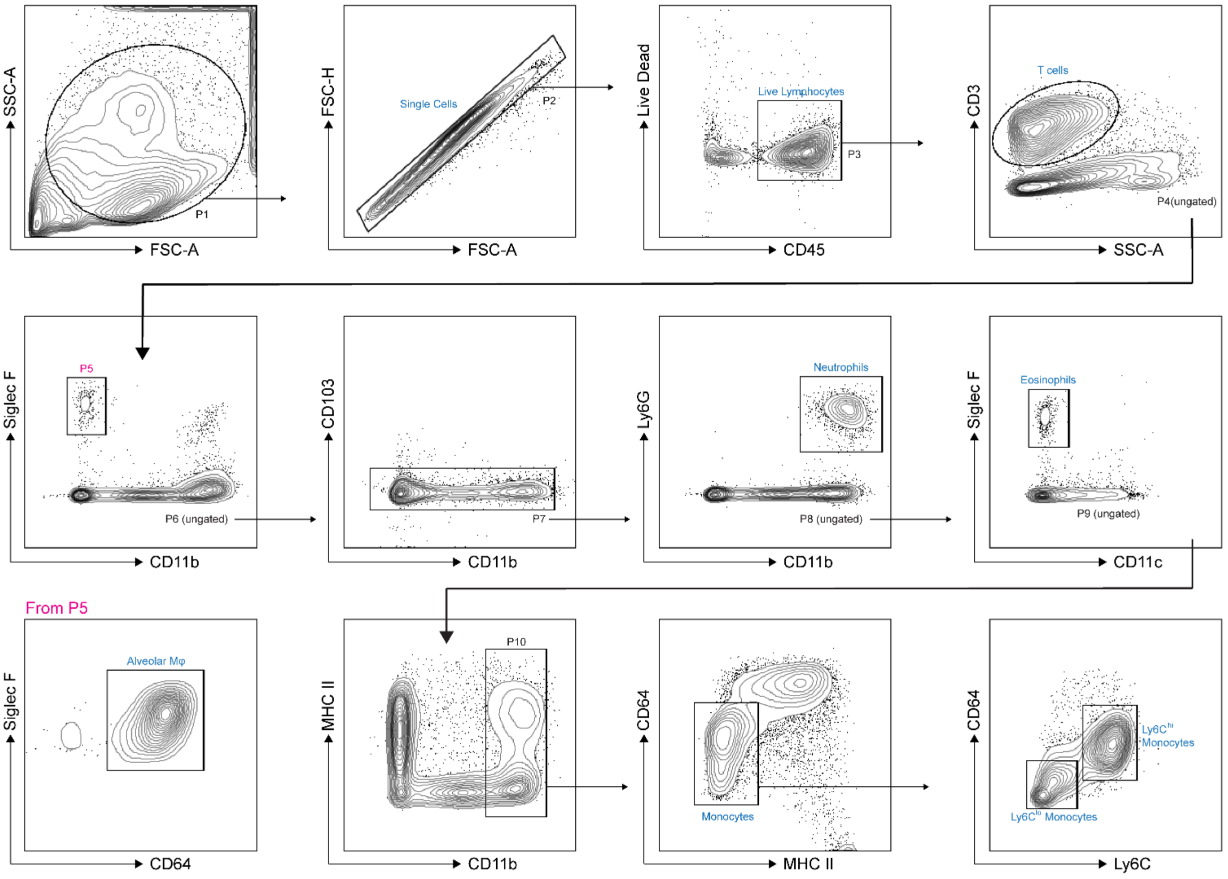


**Supplemental Figure 1. Flow cytometry gating strategy for lung digest staining.**

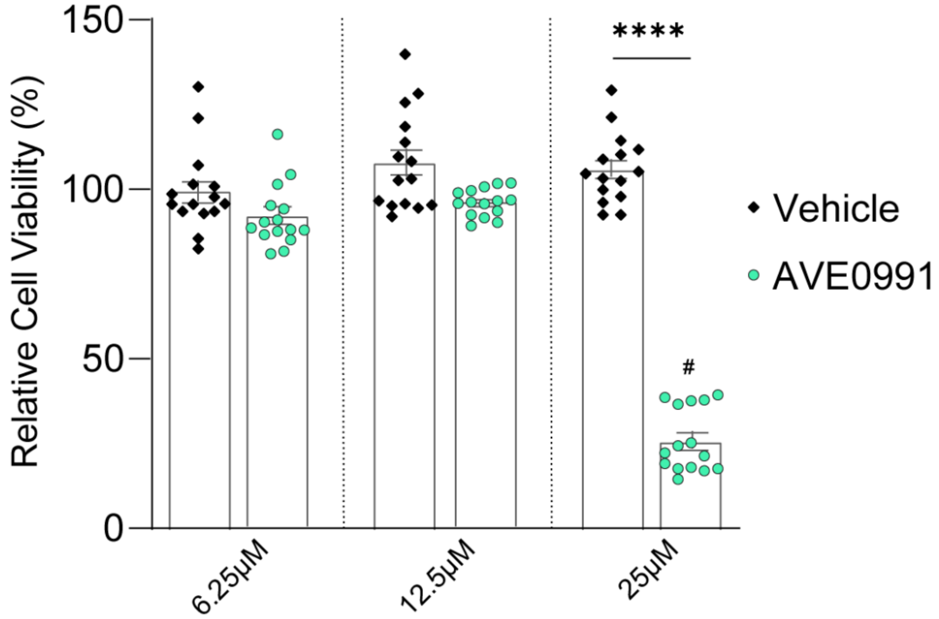


**Supplemental Figure 2. AVE0991 is cytotoxic at 25µM in Calu-3 cells.** Calu-3 cells were treated with AVE0991 (6.25, 12.5, or 25 µM), vehicle (DMSO), or media only for 24 hours. ATP levels were measured using the ATPlite luminescence assay. Cell viability is shown as percentage relative to media-only control. Normality was assessed by Shapiro-Wilk test. Statistical analysis was performed using Kruskal-Wallis test with Dunn's multiple comparisons. Data are pooled from three independent experiments and presented as mean ± SEM. ****: $p<0.0001$. # indicates $P<0.05$ vs. media-only control.

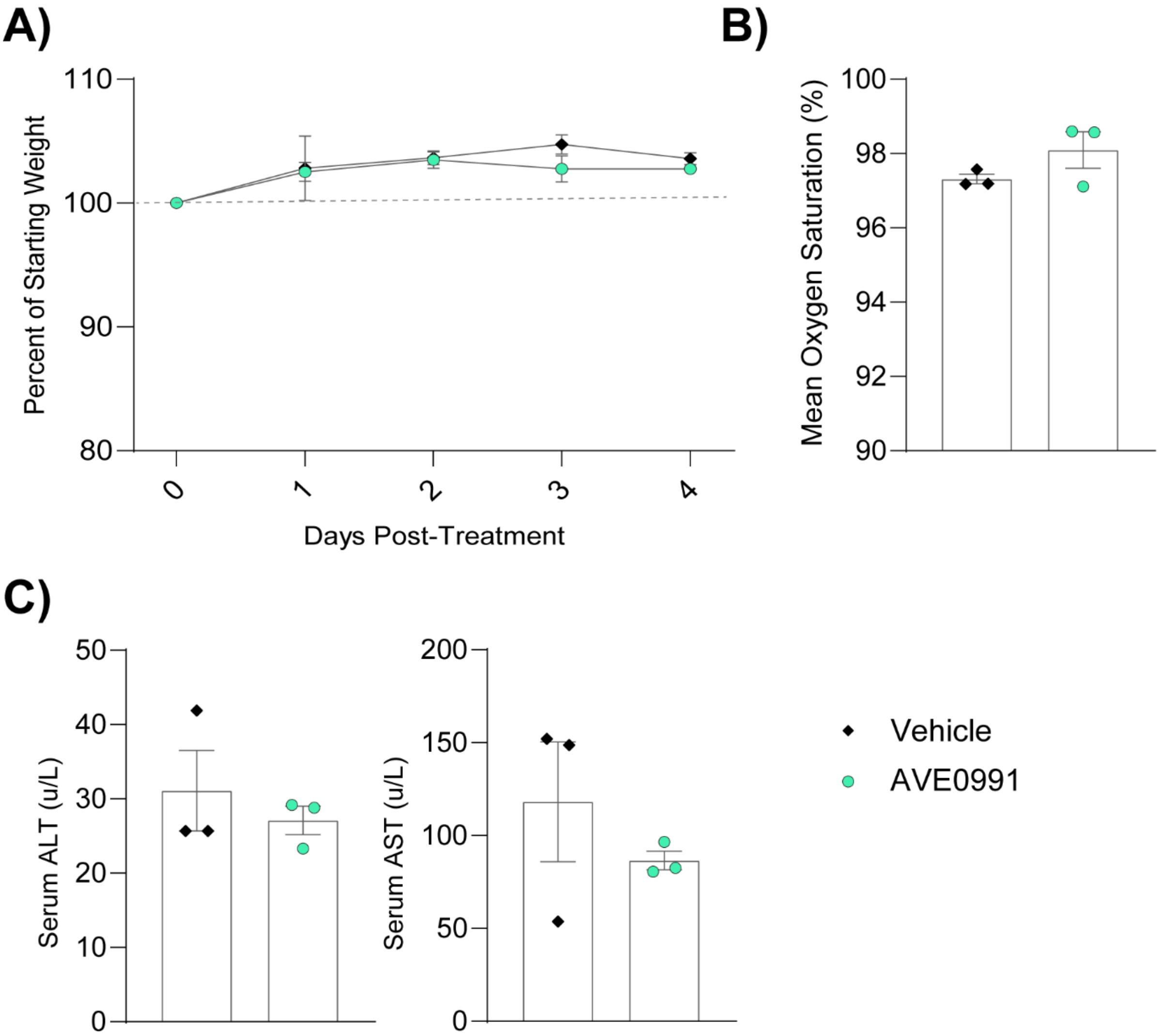


**Supplemental Figure 3. AVE0991 is not toxic in mice at 10 mg/kg oral dose.**
Male C57BL/6J mice (n=3 per group; 13-weeks old) were treated with AVE0991 (10 mg/kg) or vehicle (DMSO) via oral gavage twice daily (12-hour intervals) starting on day 0 and continuing through 4 days post-treatment. (A) Body weight shown as percentage of starting weight at treatment initiation. (B) Mean oxygen saturation. (C) Serum toxicity biomarkers. For B–C, each data point represents an individual mouse.

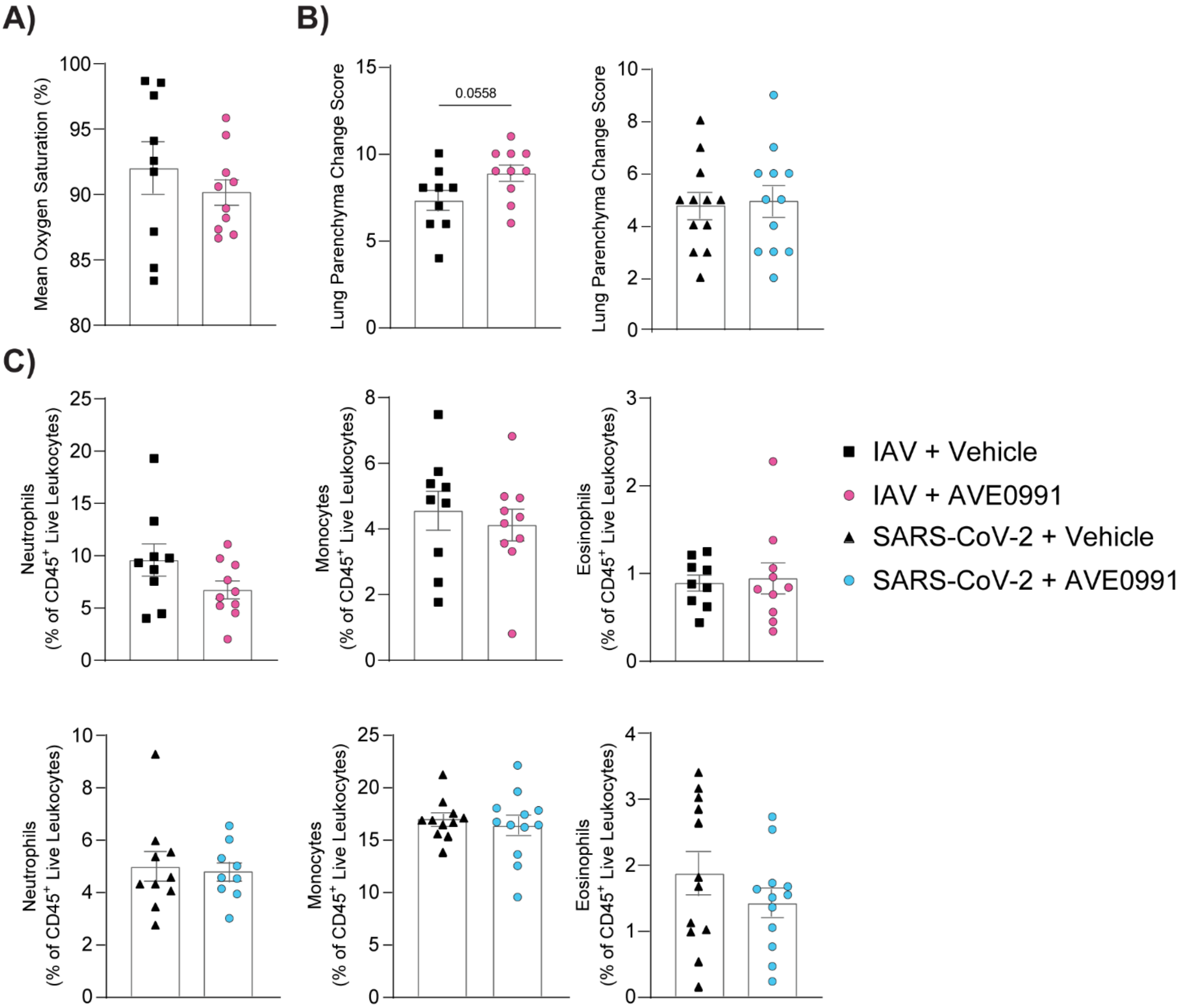


**Supplemental Figure 4. Lung characteristics of AVE0991-treated IAV- and SARS-CoV-2-infected mice.** This figure continues the data presented in Figure 2.2. Each point represents an individual mouse. (A) Mean oxygen saturation (%) in AVE0991- or vehicle-treated IAV-infected mice. (B) Lung parenchyma changes scores from histological analysis of IAV- and SARS-CoV-2-infected mice. (C) Immune cell populations in lung digests from IAV- and SARS-CoV-2-infected mice.

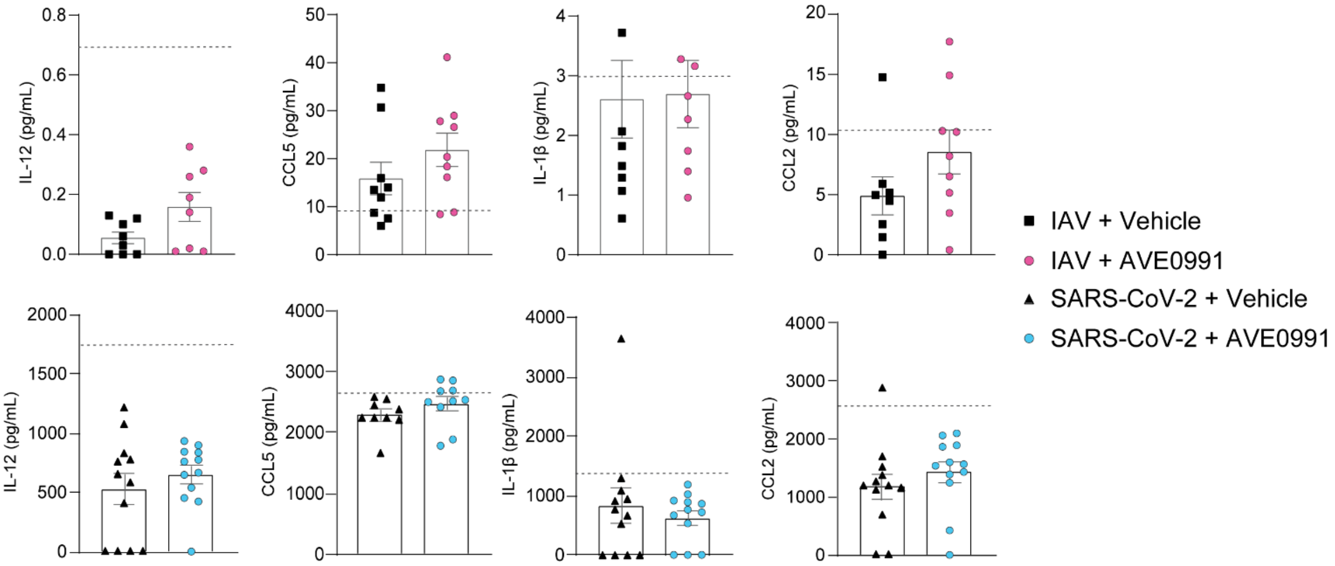


**Supplemental Figure 5. Serum cytokine levels in AVE0991-treated IAV- and SARS-CoV-2-infected mice.** This figure continues the data presented in Figure 2. Each point represents an individual mouse. The dotted line indicates the assay's limit of detection.

### Supplemental Tables

**Supplemental Table 1. Primer sequences for qPCR**

| Target | Forward or Reverse | Primer Sequence |
|---|---|---|
| **GAPDH** | Forward | 5’-CGAGATCCCTCCAAAATCAA-3’ |
| | Reverse | 5’-TTCACACCCATGACGAACAT-3’ |
| **IL-6** | Forward | 5’- CTCAGCCCTGAGAAAGGAGACAT-3’ |
| | Reverse | 5’- TCAGCCATCTTTGGAAGGTTCA-3’ |
| **TNF-α** | Forward | 5’- AGCCCATGTTGTAGCAAACC-3’ |
| | Reverse | 5’- TGAGGTACAGGCCCTCTGAT-3’ |

**Supplemental Table 2. Lung digest antibody panel**

| Marker | Company & Cat. number | Antibody | Clone | Fluorophore | Concentration per sample |
|---|---|---|---|---|---|
| **CD45** | BD 561037 | Rat LOU | 30-F11 | APC-Cy7 | 2μg/ml |
| **CD11b** | BD 563553 | Rat DA | M1/70 | BUV395 | 2μg/ml |
| **Ly6C** | BD 563011 | Rat IgM, κ | AL-21 | BV605 | 1μg/ml |
| **Ly6G** | BD 551461 | Rat LEW | 1A8 | PE | 0.25μg/ml |
| **Siglec F** | BD 562757 | Rat LOU | E50-2440 | PE-CF594 | 1μg/ml |
| **CD103** | BD 562772 | Rat LOU | M290 | APC | 2μg/ml |
| **CD64** | Biolegend 139301 | Mouse IgG1, κ | X54-5/7.1 | BV421 | 10μg/ml |
| **MHCII** | BD 74894 | Rat BN x LEW IgG2b, κ | M5/114.15.2 | BV786 | 0.25μg/ml |
| **CD11c** | BD 612796 | Hamster IgG1, λ2 | HL3 | BUV737 | 2μg/ml |
| **CD3** | Biolegend 100203 | Rat IgG2b, κ | 17A2 | FITC | 0.5μg/ml |
